# Systematic identification of regulators of antibody-drug conjugate toxicity using CRISPR-Cas9 screens

**DOI:** 10.1101/557454

**Authors:** C. Kimberly Tsui, Robyn M. Barfield, Curt R. Fischer, David W. Morgens, Amy Li, Benjamin A. H. Smith, Carolyn R. Bertozzi, David Rabuka, Michael C. Bassik

## Abstract

Antibody-drug conjugates (ADCs) selectively deliver highly toxic chemotherapeutic agents to target antigen-expressing cells and have become an important cancer treatment in recent years. However, the molecular mechanisms by which ADCs are internalized and activated within cells remain unclear. Here we use CRISPR-Cas9 screens to identify genes that control the toxicity of ADCs. Our results demonstrate critical roles for a range of known and novel endolysosomal trafficking regulators in ADC toxicity. We identify and characterize C18orf8/RMC1 as a regulator of ADC toxicity through its role in endosomal maturation. Through comparative analysis of CRISPR screens with ADCs bearing a noncleavable linker versus a cleavable valine-citrulline (VC) linker, we show that a subset of late endosomal and lysosomal regulators are selectively essential for toxicity of noncleavable linker ADCs. We further show that cleavable VC linkers are rapidly processed upon internalization and therefore surprisingly appear to bypass the requirement of lysosomal delivery. Lastly, we show that inhibition of sialic acid biosynthesis sensitizes cells to ADC treatment by increasing the rate of ADC internalization. This sensitization was observed using several ADCs targeting different antigens in diverse cancer cell types, including the FDA-approved ADC trastuzumab emtansine (T-DM1) in Her2-positive breast cancer cells. Together, these results reveal novel regulators of endolysosomal trafficking, provide important insights to guide future ADC design, and identify candidate combination therapy targets as well as potential mechanisms of ADC resistance.

## Introduction

Antibody-drug conjugates (ADCs) are an emerging class of targeted cancer therapeutics with immense promise and demonstrated clinical success^1–6^. ADCs combine monoclonal antibodies with highly toxic small molecules to selectively deliver chemotherapeutic agents to antigen-expressing tumor cells. Currently, four ADCs have been approved by the FDA since 2013 and more than 80 distinct ADCs are currently being tested in clinical trials for a range of cancers^6^. Despite intense clinical interest, the mechanisms by which ADCs enter and kill the cell – a process comprising many steps including internalization, intracellular trafficking, catalytic processing, and lysosomal escape – remain incompletely understood.

Our current understanding of ADC uptake and trafficking is largely based on classical studies of endogenous ligands and their receptors. ADCs are thought to bind their target antigen on the cell surface, undergo internalization, and traffic to the lysosome. In the lysosome, ADCs are processed by proteolytic and other enzymes that cleave the conjugated drug from the antibody, triggering “payload” release and toxicity. However, antibody binding may alter receptor internalization and trafficking pathways, leading to a decrease in delivery to the lysosome and increased delivery to other cellular compartments^7–9^, resulting in suboptimal killing. Altered trafficking routes have also been implicated in resistance to ADC treatments^9–11^, highlighting the need for a comprehensive understanding of how ADC trafficking is controlled.

The chemical linkage between the antibody and the conjugate drug is a critical feature that ensures stability of ADCs in circulation while allowing their toxic payload to be released within target cells. Linkers used for ADCs are often classified as either “cleavable” or “noncleavable”, and both designs are being used in the clinic^2^. Cleavable linkers have been engineered to be sensitive to enzymes, acidic, or reductive conditions, thereby releasing the ADC payload upon exposure to these intracellular stimuli^2^. However, some cleavable linkers have been associated with non-specific, extracellular release and off-target killing^12–14^. In contrast, noncleavable linkers are more stable in circulation as they require antibody degradation to become cytotoxic^2, 12^. A significant complication for noncleavable linkers is that their payload remains attached to the linker and the conjugated amino acids, assemblies that tend to be membrane-impermeable and require the activity of lysosomal transporters to escape into the cytosol^4^.

The optimal linker type remains an open question, as existing clinical and preclinical data have been decidedly mixed. For example, preclinical studies of ado-trastuzumab emtansine (T-DM1/ Kadcyla®) showed that the noncleavable thioether linker (SMCC) was more effective than cleavable versions *in vivo*^15^. However, noncleavable linkers are ineffective when used in targeting certain cancer antigens^16^. While further trials and optimization could improve linker efficacy, it is likely that these results depend on intracellular trafficking of the ADC and the genetic landscape of the tumors.

Here, we use genome-wide CRISPR-knock out screens to identify modulators of ADC toxicity in an unbiased fashion, and secondary screens to identify genes with differential roles in processing of ADCs with noncleavable linkers versus cleavable valine-citrulline (VC) linkers. We uncovered many known and novel endolysosomal regulators that influence ADC trafficking and processing, revealing potential resistance mechanisms. We identified C18orf8/RMC1 as a novel modulator of ADC toxicity and further characterized its role in general endolysosomal trafficking. Lastly, we found that inhibition of sialic acid biosynthesis sensitizes cells to anti-CD22 ADC treatment by boosting rate of ADC internalization, a finding which generalized to the FDA-approved Her2-targeting ADC trastuzumab emtansine (T-DM1). Together, our results elucidate mechanisms controlling ADC trafficking and toxicity, and provide insight for future ADC development.

## Results

### Genome-wide CRISPR screen uncovers diverse endolysosomal regulators of ADC toxicity

To identify genes regulating ADC toxicity, we conducted a genome-wide CRISPR-deletion screen using a potent CD22-targeting ADC (anti-CD22-Asparagine-PEG2-maytansine, hereinafter referred to as anti-CD22-maytansine). The ADC was synthesized using a site-specific conjugation technology based upon the aldehyde tag ^17^ and HIPS chemistry ^18^ to attach a microtubule inhibitor (maytansine^19^) payload coupled through a noncleavable linker to the antibody heavy chain C-terminus (Supplementary Fig. 1a). Maytansine is a highly toxic microtubule inhibitor used in many ADCs^2^. We first tested that the anti-CD22-maytansine ADC inhibits growth of a CD22-positive B-lymphocyte Burkitt’s lymphoma cell line (Ramos) in a dose- and antibody-dependent manner (Supplementary Fig. 1b). Then we lentivirally transduced a previously validated sgRNA knock-out library into Cas9-expressing Ramos cells targeting all protein coding genes with 10 sgRNAs/gene and ∼10,000 negative controls^20^. We then split the cells into two conditions in duplicate, one treated with 2 nM anti-CD22-maytansine for three rounds treatment (each killing ∼50% of cells) and the other left untreated. During the 2-week course of the screen, cells with sgRNAs targeting genes required for ADC toxicity are enriched in the ADC treated conditions relative to the untreated conditions while cells with sgRNAs targeting genes that when knocked out sensitize cells to ADC are depleted. The proportion of each sgRNA in the two conditions was measured by deep sequencing and significant regulators of ADC toxicity were identified using casTLE (Fig. 1a)^21^.

**Figure 1:**
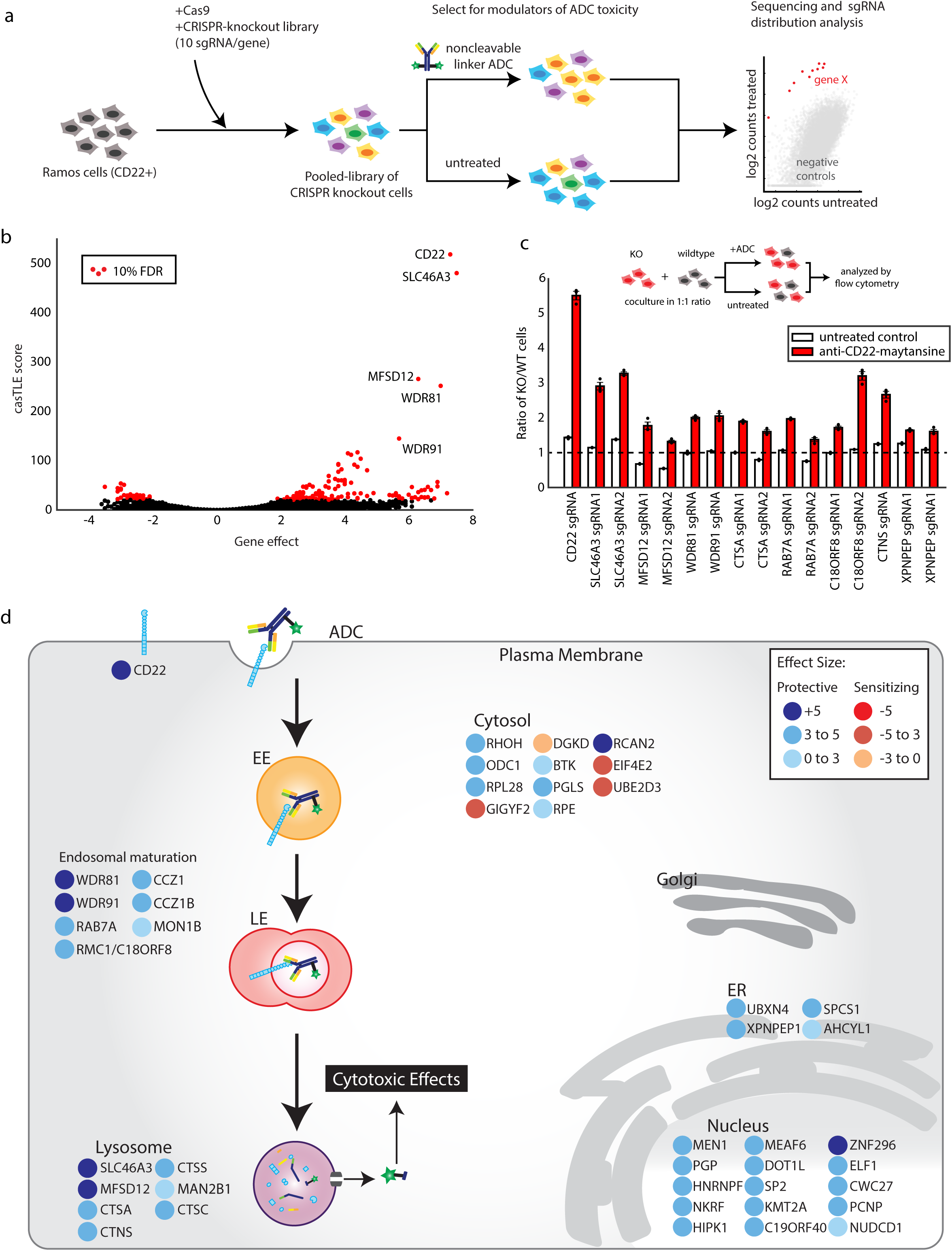
Genome-wide CRISPR screen uncovers diverse endolysosomal regulators of ADC toxicity. a. Schematic for the genome-wide screen. Ramos cells expressing Cas9 were infected with a lentiviral, genome-wide sgRNA library, the population was split, and half the cells were treated with anti-CD22-maytansine ADC for three rounds, killing ∼50% cells each round. The resulting populations were subjected to deep sequencing and analysis. The screen was performed in duplicate. b. Volcano plot of all genes indicating effect and confidence scores for genome-wide ADC screen. Effect and confidence scores determined by casTLE. c. Validation of hits in Ramos cells using competitive-growth assays: Cells expressing sgRNAs for knock out (KO) of indicated genes (mCherry+) and control (mCherry-) were co-cultured in 1:1 ratio. Cells were either treated with anti-CD22-maytansine ADC or left untreated for three days. Resulting ratio of KO:control was determined using flow cytometry (n=3; error bars, ± SEM). d. Schematic for top 30 and selective endolysosomal traffi cking regulators hits in genome-wide screen, color coded by effect size (calculated by casTLE). Subcellular localization are indicated according to GO annotations.

This initial screen identified 168 known and novel regulators of ADC toxicity at a 10% false discovery rate (FDR) (Fig. 1b and 1d; see Supplementary Table 1 for full list). Top hits were then validated with individual sgRNA-expressing knockout cell lines in competitive-growth assays (Fig. 1c). Among the strongest hits was CD22, the target antigen of the ADC, as well as SLC46A3, a previously reported lysosomal transporter that is required for maytansine-conjugates with noncleavable linkers to enter the cytosol after proteolytic processing^22^. These positive controls indicated the screening strategy could robustly identify genes critical for ADC biology. Moreover, we identified a number of endolysosomal genes as modifiers of ADC toxicity, including RAB7A (a key regulator of late endolysosomal trafficking^23–25^), MON1 and CCZ1 (both activators of RAB7A activity^26–28^). The screen revealed that WDR81 and WDR91, two recently characterized regulators of endosomal maturation^29, 30^, are essential for ADC toxicity, suggesting that they also play roles in mediating ADC lysosomal delivery. Deletion of lysosomal cathepsins A, C, and S were highly protective against ADC killing; these enzymes are likely important for proteolytic degradation of the antibody. We also identified many nuclear genes such as chromatin modifiers MEN1^31, 32^ and DOT1L^33^, which may be required for the anti-mitotic activity of maytansine or are indirectly influencing ADC trafficking and processing.

### Targeted screens reveal critical role of lysosomal delivery in toxicity of noncleavable linker ADC but not cleavable linker ADC

We next aimed to evaluate whether the genetic modifiers identified in our genome-wide screen play roles in ADC trafficking, processing, or the anti-mitotic activity of maytansine. To do this, we performed parallel high cell coverage targeted screens using an ADC with a noncleavable linker (anti-CD22-maytansine used above), an ADC with a cleavable linker (anti-CD22-Glutamate-PEG2-valine-citrulline-maytansine, hereinafter referred to as anti-CD22-VC-maytansine, see supplementary Fig. 1b for structure), and free maytansine. These three treatments were screened in parallel in Ramos cells (Fig. 2a), using a targeted sublibrary of sgRNAs targeting (1) the hits identified in the genome-widescreen, (2) manually curated endolysosomal regulators, and (3) lysosomal-localized genes^34^ (Fig. 2a, see Supplementary Table 5 for list of genes and sgRNAs). In contrast to noncleavable linkers, valine-citrulline (VC) linkers are designed to be cleaved by the cysteine protease cathepsin B^35, 36^ (Supplementary Fig. 1c)., but were recently shown to be also sensitive to other cathepsins^37^. Proteolytic processing of anti-CD22-VC-maytansine releases a maytansine payload that is expected to freely diffuse across endosomal membranes. Similar to the genome-wide screens, screens for each condition were performed in duplicate and either left untreated, or treated for three rounds of selection with ADCs or free maytansine at concentrations that killed ∼50% of the cells. Hits were identified using casTLE^21^ by comparing each treated condition to the untreated control (see Supplementary Table 2 for complete list of hits).

**Figure 2:**
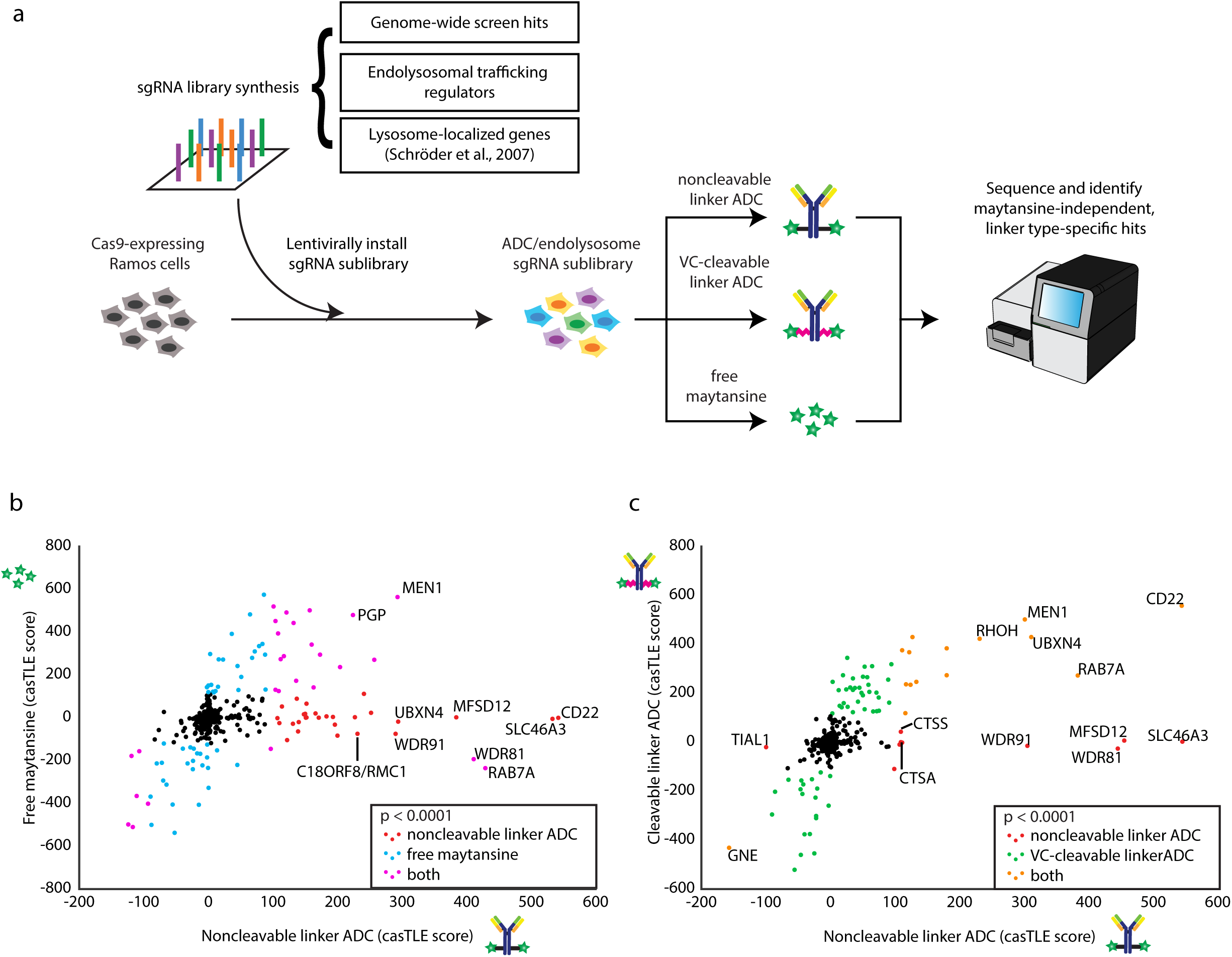
A subset of endolysosomal traffi cking regulators are critical only for toxicity of the noncleavable linker ADC. a. Schematic for sublibrary targeted screen using anti-CD22-maytansine (noncleavable), anti-CD22-VC-maytansine (cleavable), and free maytansine. The targeted sublibrary was designed, synthesized, and lentivirally installed into Cas9-expressing Ramos cells. The Ramos cells were either treated with (1) anti-CD22-maytansine, (2) anti-CD22-VC-maytansine, (3) free maytansine, or left untreated. The resulting populations were subjected to deep sequencing and analysis. The screens were performed in duplicate and at 1000x coverage. b. Comparative analysis of results from anti-CD22-maytansine (noncleavable) and free maytansine sublibrary screens. Signed casTLE scores are reported. c. Comparative analysis of results from anti-CD22-maytansine (noncleavable) and anti-CD22-VC-maytansine (cleavable) sublibrary screens. Signed casTLE scores are reported.

To distinguish genes involved in the activity of maytansine from those influencing ADC trafficking and processing, we compared hits identified in the free maytansine condition to hits obtained using noncleavable anti-CD22-maytansine (Fig. 2b, supplementary Fig. 2). Genes identified in both conditions, which includes most nuclear genes hits, are likely to mediate the killing activity of maytansine. On the other hand, hits unique to the ADC are likely to influence mechanisms of ADC trafficking, processing, or lysosomal escape. These include the target-antigen CD22, intracellular trafficking regulators (RAB7A, WDR81, and WDR91), lysosomal cathepsins (CTSS and CTSA), and lysosomal transporters (SLC46A3 and MFSD12), highlighting the importance of lysosomal delivery and processing in ADC toxicity.

To investigate how particular linkers influence ADC toxicity, we compared the hits obtained using the noncleavable linker ADC vs. the cleavable VC linker ADC (Fig 2C, supplementary Fig. 2). As expected, CD22 along with hits identified in the free maytansine condition were common hits for both ADCs. We did not identify specific proteases that are essential for toxicity of anti-CD22-VC-maytansine, likely due to redundancy as the VC linker can be cleaved by multiple cathepsins^37^. Strikingly, a subset of endolysosomal trafficking regulators were identified only to be critical for toxicity of the noncleavable linker ADC but were dispensable for the cleavable linker ADC, suggesting that lysosomal delivery may not be required for processing of VC linkers.

### Lysosomal delivery is not required for processing of cleavable valine-citrulline-linked ADCs

Next, we sought to elucidate the mechanistic differences of proteolytic processing and payload release of noncleavable linker ADC and cleavable linker ADC. We first validated that the subset of endosomal and lysosomal genes identified in our targeted screen are indeed specific to the noncleavable linker ADC (SLC46A3, MFSD12, WDR81, WDR91, CTSA, CTSS and C18orf8/RMC1) and have little effect on toxicity of the VC cleavable linker ADC in competitive growth assays (Fig. 3a).

**Figure 3:**
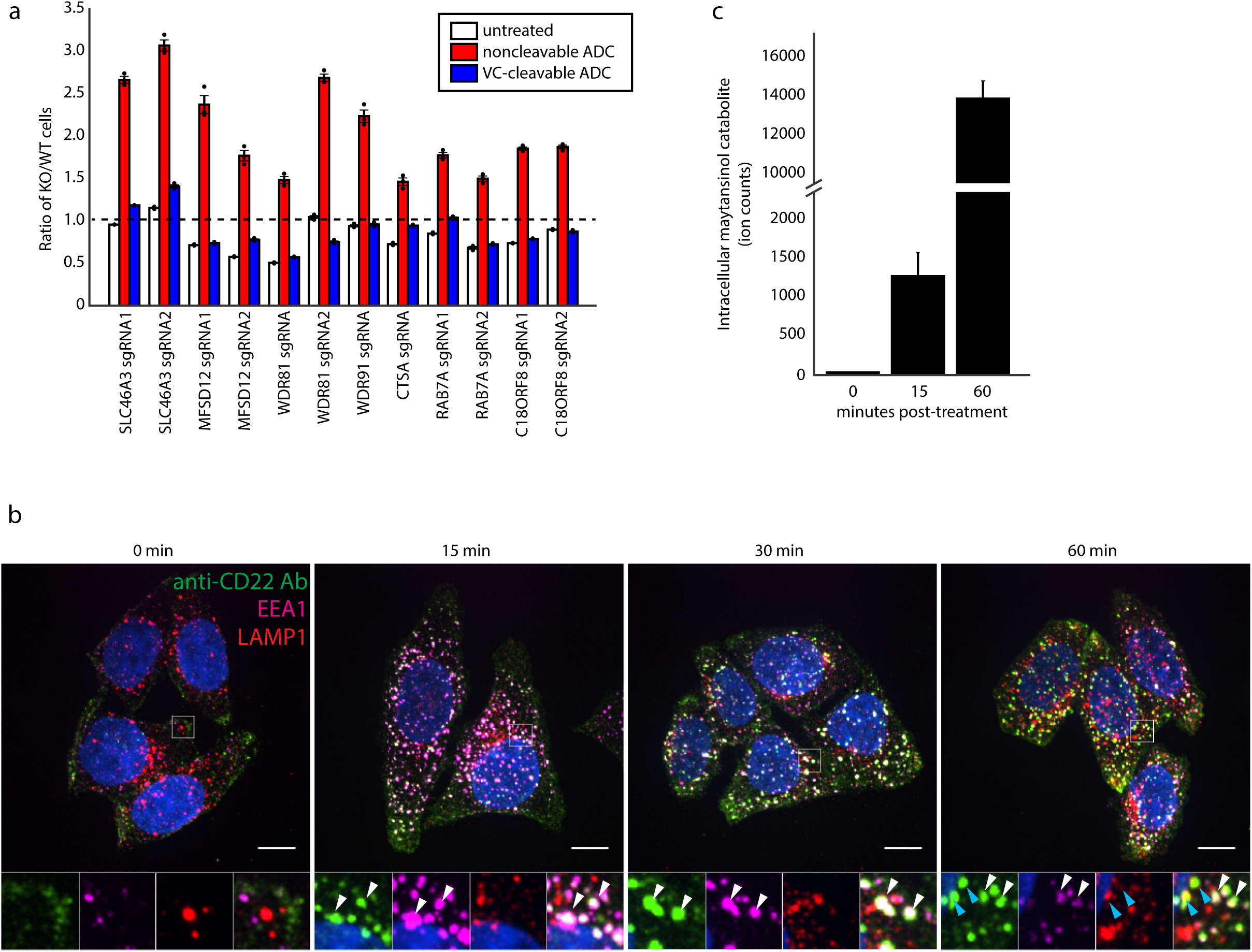
Lysosomal delivery is not required for processing of VC cleavable linkers. a. Validation of hits in Ramos cells using competitive-growth assays: Cells expressing sgRNAs for KO of indicated genes (mCherry+) and control (mCherry-) were co-cultured in 1:1 ratio. Cells were either treated with anti-CD22-maytansine, anti-CD22-VC-maytansine (blue) or left untreated (white) for three days. Resulting ratio of KO:control was determined using flow cytometry (n=3; error bars, ± SEM). b. Internalization of anti-CD22 antibody conjugated to Alexa Fluor 488 (anti-CD22-AF488) as monitored by immunofluorescence microscopy in HeLa cells. Cells were incubated on ice with anti-CD22-AF488 for 20 minutes. Cells were then washed once with cold PBS and internalization was initiated by pre-warmed media. Cells were washed and fixed at indicated time points post-internalization initiation. White arrows indicate colocalization of anti-CD22 antibody and early endosomal marker EEA1; blue arrows indicate colocalization of anti-CD22 antibody and late endosomal/lysosomal marker LAMP1. c. Maytansinol release from anti-CD22-VC-maytansine in CD22-expressing HeLa cells. 10 nM anti-CD22-VC-maytansine was added to cells and reactions were quenched at indicated time points. Level of intracellular maytansinol catabolite was determined by LC/MS-MS; see methods for detailed extraction and detection protocol (n=3, normalized by cell number and internal standard MMAE; error bars, ± SEM).

Because we find that toxicity of ADCs with cleavable VC linkers is not dependent on genes regulating endosomal maturation or specific lysosomal proteases, we hypothesize that the VC cleavable linker can be processed in earlier endosomal compartments, and may be able to bypass defects in lysosomal delivery. To monitor ADC intracellular trafficking, we stably expressed CD22 in HeLa cells (Supplementary Fig. 3a) and monitored trafficking of a fluorescently-labeled CD22 antibody. We found the ADC colocalized with an early endosomal marker EEA1 at 15 minutes after treatment and began to colocalize with lysosomal marker LAMP1 in approximately 1 hour (Fig. 3b). To detect cleavage and release of the payload, we used liquid chromatography coupled with tandem mass spectrometry (LC-MS/MS) to detect intracellular levels of the maytansinol cleavage product^38^ (Supplementary Fig. 3b and 3c). The cleaved maytansinol product began to appear as early as 15 minutes post-treatment (Fig. 3c and Supplementary Fig. 3d, coinciding with the accumulation of anti-CD22 antibody in early endosomes as assayed by microscopy, but prior to accumulation in the lysosome (Fig3b and 3c). The maytansinol product continued to accumulate over the course of the experiment (Supplementary Fig. 3d). Notably, inhibition of V-ATPase using bafilomycin A1^39^ prevents maytansinol payload release, indicating that cleavage of the VC linker is dependent on proteases that require proper acidification of endosomes and lysosomes (Supplementary Fig. 3e). Together, these results highlight the differences in processing of ADCs with noncleavable linker and VC linker, and provide evidence that lysosomal delivery is not critical for toxicity of ADC with a cleavable VC linkers.

### C18orf8/RMC1 is required for endosomal trafficking and lysosomal delivery of ADCs

A top maytansine-independent hit in our screen was C18orf8/RMC1 (hereinafter referred to as RMC1), a gene recently identified as an autophagy regulator acting in concert with MON1 and CCZ1 to activate Rab7^40^. As knockout of either RAB7A or RMC1 protects cells from ADC-mediated toxicity (Fig. 3a), we hypothesized that RMC1 plays a role in endosomal maturation through regulation of RAB7A activity, and that loss of RMC1 activity would prevent lysosomal delivery of ADCs and cell death. To investigate the role of RMC1 in endolysosomal trafficking, we first examined the localization of a C-terminal GFP-fused RMC1 in HeLa cells and found it colocalizes with the lysosomal marker LAMP1 (Fig. 4a), as previously reported. Next, we knocked down RMC1 or RAB7A using CRISPR interference^41^ (CRISPRi) in HeLa cells. Loss of RMC1 or RAB7A function both led to enlarged early endosomes and lysosomes (Fig. 4b), accompanied by increased EEA1 and LAMP1 staining (Supplementary Fig. 4a). To determine whether these changes in morphology affect cargo transport to the lysosomes, we monitored trafficking of epidermal growth factor (EGF), a canonical endocytic cargo that traffics to the lysosome using confocal microscopy. Indeed, in RMC1-knock down cells EGF trafficking to the lysosome is markedly delayed and EGF accumulated in early endosomes (Fig. 4c). Consistently, lysosomal degradation of EGFR upon EGF treatment was also impaired (Fig. 4d), confirming the role of RMC1 in lysosomal delivery of endosomal substrates. Finally, to determine how RAB7A activity influences the role of RMC1 in ADC toxicity, we tested whether overexpression of constitutively active RAB7-Q67L re-sensitizes RMC1-deleted cells to noncleavable linker ADCs. Indeed, RMC1-deleted cells overexpressing RAB7-Q67L were more sensitive to ADC treatment than cells expressing GFP control (Fig. 4f). Together, these results support the hypothesis that RMC1 is essential for ADC delivery and toxicity in a RAB7-dependent manner, and more generally demonstrate a role for RMC1 in delivery of endocytic substrates to the lysosome.

**Figure 4:**
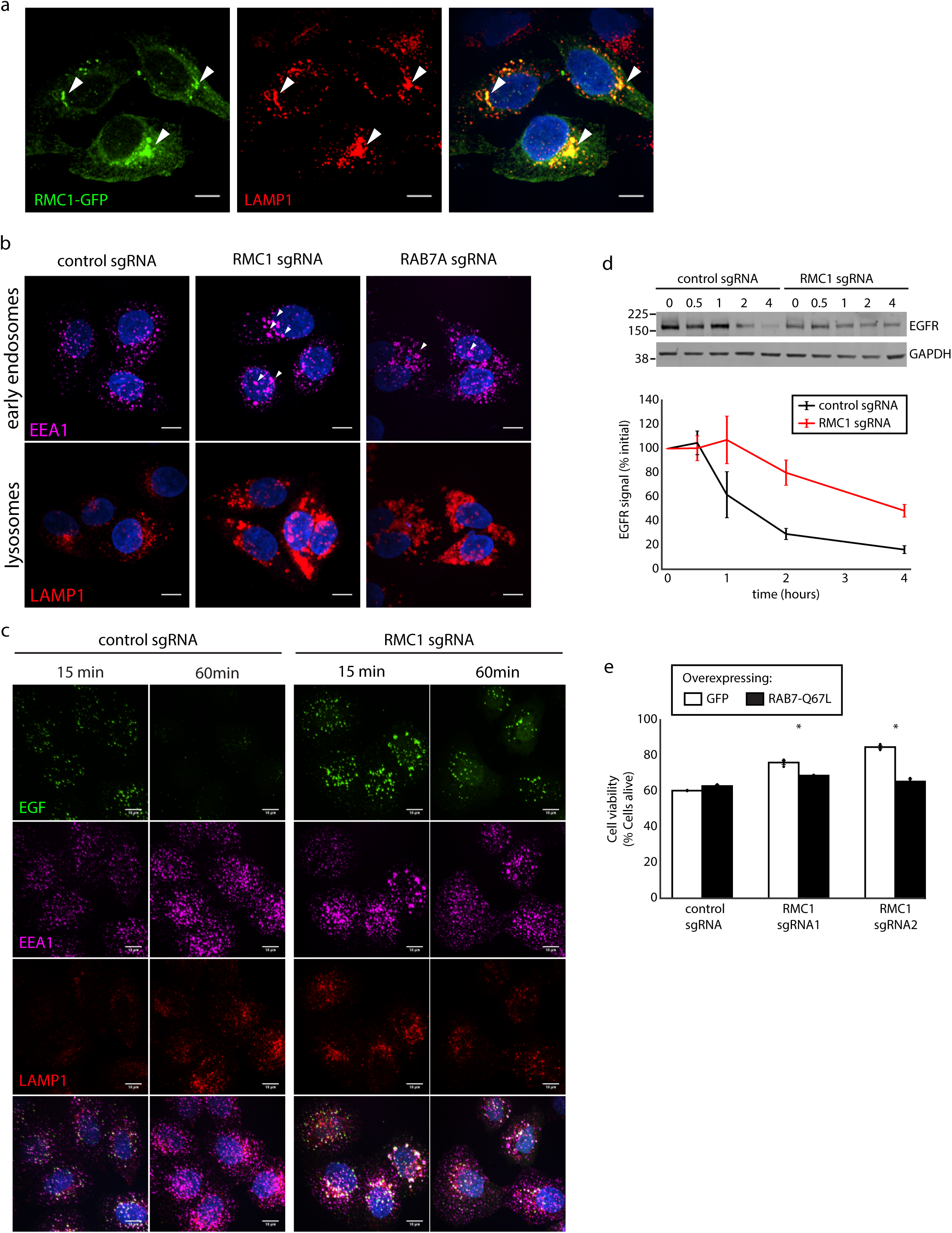
C18orf8/RMC1 is required for endosomal traffi cking and lysosomal delivery of ADCs. a. C-terminal GFP-tagged RMC1 localize to LAMP1-positive late endosome/lysosomes, visualized by immunofluorescence and confocal microscopy. White arrows indicate sites of colocalization. Scale bars, 10 µm. b. CRISPRi-mediated knock down of RMC1 or RAB7A in HeLa cells result in enlarged endosomes (EEA1) and lysosomes (LAMP1). Scale bars, 10 µm. c. Trafficking of EGF in RMC1-knock down HeLa cells. Cells were serum-starved for 16 hours, then were placed on ice and treated with EGF for 10 minutes. Cells were then washed with once with cold PBS and internalization was initiated by pre-warmed media. Cells were fixed at indicated time and visualized by immunofluorescence and confocal microscopy. Scale bars, 10 µm. d. EGFR degradation in RMC1-knock down HeLa cells. Cells were treated with EGF for 10 min, and then EGFR levels were examined at indicated time point by Western blot. EGFR levels were quantified with ImageJ software (n=3; error bars, ± SEM). e. Overexpression of constitutively active RAB7A-Q67L restores sensitivity towards anti-CD22-maytansine (noncleavable) in RMC1-knockout Ramos cells. Cells were treated with 1 nM anti-CD22 noncleavable linker ADC and cell viability was analyzed after three days of ADC treatment by flow cytometry. Viability was as determined by forward and side scatter of live gating of Ramos cells (n=3; Two-tailed t test, *P<0.05; error bars, ± SEM).

### Inhibition of sialic acid synthesis sensitizes cells to CD22 ADC toxicity

Genes whose deletion sensitizes cancer cells to ADCs could potentially serve as targets for combination therapies to increase ADC efficacy in the clinic. Although the genome-wide and follow-up screens identified many genes and pathways required for ADC toxicity, we identified relatively few genes whose deletion robustly sensitized cells to ADC treatment. We hypothesized that this was due to either insufficient cell coverage (∼1000 cells per sgRNA) or the high dose of ADC used in the screens, both of which could limit the dynamic range of sensitization. We thus repeated the screen at a higher cell coverage (∼5000 cells per sgRNA) and with lower dose of the noncleavable linker ADC (0.5 nM) to specifically search for sensitizing targets (Fig. 5a). To prioritize identification of candidate combination therapy targets, we screened in a targeted sublibrary highly enriched in known drug targets, kinases, and phosphatases. As we were most interested in finding genes that influence ADC trafficking and processing, a free maytansine treatment was also used as a counter-screen to filter out maytansine-dependent genes. As expected, this design identified a number of strong sensitizing hits that were missed in the genome-wide screen (Supplementary Fig. 5a-c; see Supplementary Table 3 for complete list of hits).

**Figure 5:**
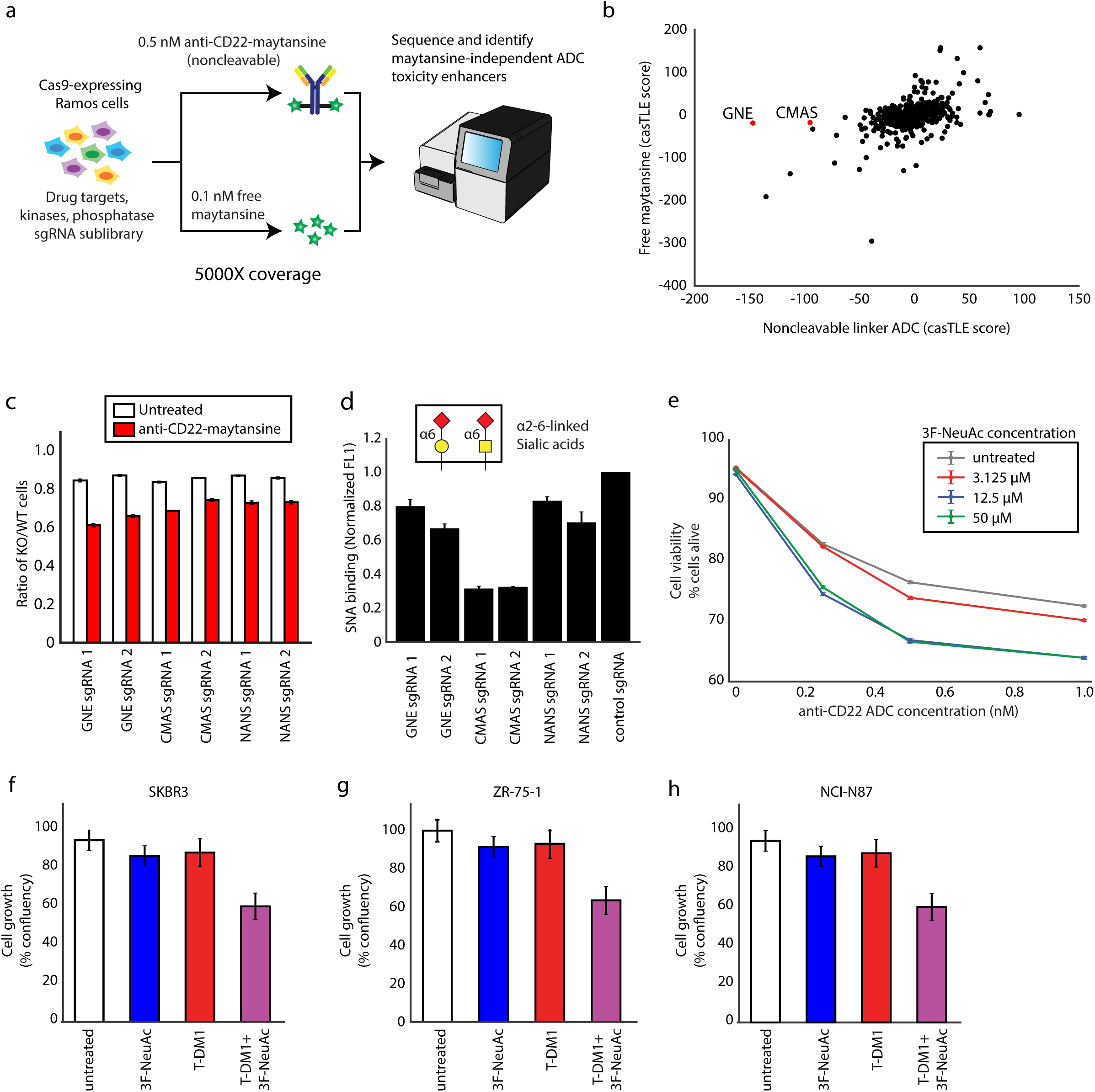
Inhibition of sialic acid synthesis sensitizes cells to CD22 ADC toxicity. a. Schematic for targeted screen for identifying sensitizing hits. Ramos cells expressing Cas9 were infected with a lentiviral, sgRNA sublibrary targeting drug targets, kinases and phosphatases. Cells were then treated with either 0.5 nM anti-CD22-maytansine, 0.1 nM free maytansine, or left untreated. The resulting populations were subjected to deep sequencing and analysis. The screen was performed in duplicate and at 5000x coverage. b. Comparative analysis of results from anti-CD22 noncleavable linker ADC (0.5 nM) and free maytansine (0.1 nM) sublibrary screens performed at 5000X coverage. c. Validation of hits in Ramos cells using competitive-growth assays. Cells expressing sgRNAs for KO of indicated genes (mCherry+) and control (mCherry-) were co-cultured in 1:1 ratio. Cells were either treated with anti-CD22 noncleavable ADC or left untreated for three days. Resulting ratio of KO:control was determined using flow cytometry (n=3; error bars, ± SEM). d. Levels of α-2,6 sialic acid on KO Ramos cells as measured by labeling with SNA, a lectin that preferentially binds to α-2,6 sialic acid residues on N- and O-glycans (schematic illustrated in figure; red diamond: sialic acid/Sia, yellow square: N-acetylglucosamine/GalNAc, yellow circle: galactose/Gal). SNA conjugated to AF488 was incubated with cells on ice for 30 minutes. Cells were then washed 3x with cold PBS and analyzed by flow cytometry (n=3; error bars, ± SEM). e. Cell viability Ramos cells treated with noncleavable ADC and 3F-NeuAc at different concentration. Cells were first treated with 3F-NeuAc at indicated concentrations for 48 hours, then ADCs at indicated concentrations were added. Cell viability, determined by forward and side scatter gating for live cells, was assayed 72 hours post-ADC addition by flow cytometry (n=3; error bars, ± SEM). f-h: Cell growth (normalized confluency %) of SKBR3 (f), ZR-75-1 (g), and NCI-N87 (h) treated with 100 µM 3F-NeuAc, 2 nM T-DM1, or combination of both. Cells were pre-treated with 100 µM 3F-NeuAc for 48 hours, followed by 72-hour incubation with 2 nM T-DM1. Confluency was determined by IncuCyte s3 Live cells analysis system and normalized to maximum confluency of the untreated condition at the end of the 5-day experiment (n=3; error bars, ± SEM).

Interestingly, the top two maytansine-independent sensitizing hits, GNE and CMAS, are both members of the sialic acid biosynthesis pathway (Fig. 5b and Supplementary Fig. 5d). Using competitive growth assays, we validated that disruption of sialylation by knocking out GNE, CMAS, or NANS (an additional pathway member not represented in the sublibrary) sensitizes Ramos cells to the ADC treatment (Fig. 5c). In addition, chemical inhibition of sialylation with a fluorinated sialic acid analog, peracetylated 3Fax-NeuAc^42^ (3F-NeuAc) also sensitized Ramos cells to ADC killing (Fig. 5e). We confirmed the depletion of cell surface sialic acids in GNE, CMAS and NANS-deleted cell (Fig. 5d), and also after treatment with 3F-NeuAc, by labelling with *Sambucus nigra* agglutinin (SNA), a lectin that preferentially binds to sialic acid attached to galactose through an α-2,6 linkage^42^(Supplementary Fig. 5e). Notably, the knockout cells are also more sensitive to treatment with the cleavable linker ADC anti-CD22-VC-maytansine (Supplementary Fig. 5f), suggesting that sensitization by inhibiting sialic acid synthesis may be broadly useful for ADCs with both non-cleavable linkers and cleavable VC linkers.

To investigate whether the sensitizing effects of sialic acid depletion can be generalized to ADCs with different targets and in different cell types, we tested whether pre-treatment with 3F-NeuAc also sensitizes Her2-positive cancer cell lines to Her2-targeted ADC ado trastuzumab emtansine (T-DM1). The combination of 3F-NeuAc and T-DM1 was more toxic than ADC alone in the Her2-positive breast cancer lines SKBR3 and ZR-75-1 (Fig. 5f and 5g), and the gastric carcinoma line NCI-C87 (Fig. 5h), indicating that the strategy of combining 3F-NeuAc and ADC may be useful for multiple cancer antigen targets.

### Depletion of sialic acid enhances ADC internalization rate

To identify which steps of ADC internalization and processing are modulated by sialylation, we considered a number of models by which the cells could be sensitized. Inhibition of sialic acid synthesis may: (1) increase ADC target antigen cell surface expression and/or ADC binding, (2) enhance complement-mediated killing activity, or (3) increase the rate of ADC internalization. First, we measured CD22 expression and antibody accessibility in Ramos cells, but 3F-NeuAc treatment did not alter cell surface binding of anti-CD22 ADCs or whole cell CD22 protein levels (Supplementary Fig. 6a and 6b). Second, we tested whether the increased killing is dependent on complement activity, but the sensitizing effect of 3F-NeuAc was not abolished in media depleted of active complement factors (Supplementary Fig. 6e). Using T-DM1, we observed similar sensitization when the Her2-positive cell lines SKBR3 and ZR-75-1 were treated with 3F-NeuAc in media depleted of active complement factors (Supplementary Fig. 6f and 6g).

Lastly, we tested was whether inhibiting sialylation alters the rate of ADC uptake. We assayed ADC internalization by pHrodo-labeled anti-CD22 ADC. Cells depleted of sialic acids via either genetic deletion of GNE (Supplementary Fig. 6c) or by 48-hour pre-treatment with 3F-NeuAc (Fig. 6a) demonstrated increased rates of ADC uptake. In addition, pre-treatment of Ramos cells with 3F-NeuAc for 48 hours increased the rate of maytansinol payload release from the CD22-targeted ADC with VC cleavable linker as measured by LC/MS-MS (Supplementary Fig. 6d). To determine whether the increased uptake rate is specific to CD22, we measured the rate of internalization for an antibody targeting a different cell surface protein, CD79b, which has also been shown to be an effective ADC target (Polson et al., 2007). As a control, we measured uptake of pHrodo-labeled dextran, which is used as a marker of general fluid phase endocytosis^43^. Interestingly, treatment of Ramos cells with 3F-NeuAc increased the rate of anti-CD79 antibody uptake (Fig. 6b) but not dextran uptake (Supplementary Fig. 6g), suggesting that inhibition of sialylation may influence antibody-mediated endocytosis specifically.

**Figure 6:**
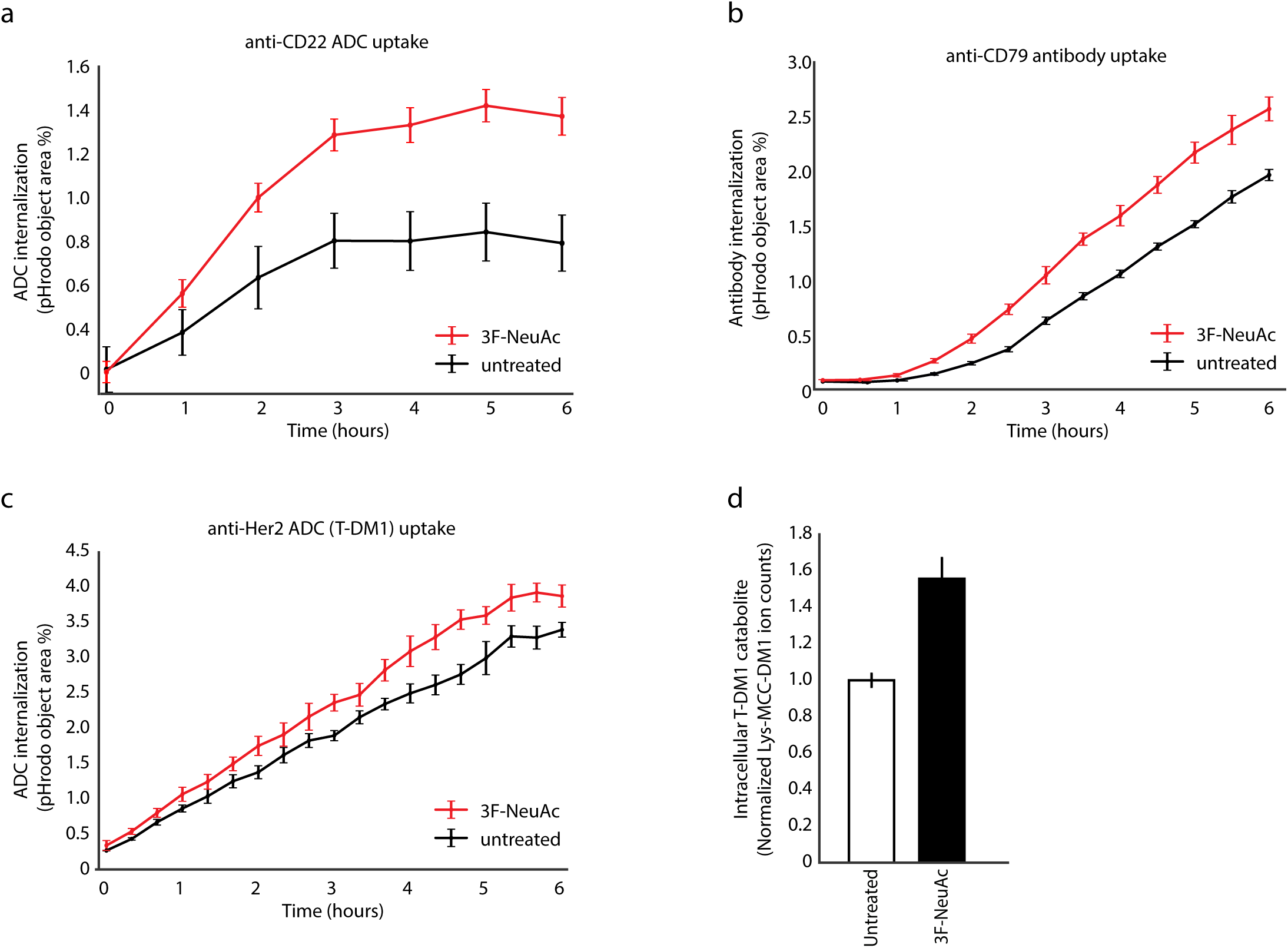
Depletion of sialic acid enhances ADC internalization rate. a. Internalization of pHrodo-labeled anti-CD22 ADC in wild type Ramos cells pre-treated with 12.5µM 3F-NeuAc for 48 hours (red) or untreated (black). pHrodo signal was measured using IncuCyte s3 Live cells analysis system and normalized to cell area (n=3; error bars, ± SEM). b. Internalization of pHrodo-labeled anti-CD79 antibody in wild type Ramos cells pre-treated with 12.5µM 3F-NeuAc for 48 hours (red) or untreated (black). pHrodo signal was measured using IncuCyte s3 Live cells analysis system and normalized to cell area (n=3; error bars, ± SEM). c. Internalization of pHrodo-labeled T-DM1 in wild type SKBR3 cells pre-treated with 100µM 3F-NeuAc for 48 hours (red) or untreated (black). pHrodo signal was measured using IncuCyte s3 Live cells analysis system and normalized to cell area (n=3; error bars, ± SEM). d. Accumulation of T-DM1 catabolite Lys-MCC-DM1 in wild type SKBR3 cells treated with 100µM 3F-NeuAc. SKBR3 cells were either pre-treated with 100µM 3F-NeuAc or left untreated for 48 hours, followed by addition of 10 nM T-DM1. After 24 hours, level of intracellular T-DM1 catabolite, Lys-MCC-DM1, was determined by LC/MS-MS; see methods for detailed extraction and detection protocol (n=3; normalized by cell number and internal standard MMAE; error bars, ± SEM).

Finally, to determine whether the effects of 3F-NeuAc on ADC internalization rate can be generalized to other ADCs and cancer cell types, we measured the rate of T-DM1 uptake and payload release using LC/MS-MS^44^ in SKBR3 cells. Indeed, T-DM1 internalization rate is enhanced in 3F-NeuAc pre-treated SKBR3 cells (Fig. 6c). In addition, increased accumulation of T-DM1’s catabolite Lys-MCC-DM1 is also observed in 3F-NeuAc pre-treated SKBR3 cells (Fig. 6d). Together, these results show that inhibition of sialylation is a generalizable strategy for increasing ADC uptake, and provide rationale for combining ADCs and inhibition of sialic acid biosynthesis to enhance the efficacy of ADC treatment.

## Discussion

Here, we used CRISPR-knockout screens to evaluate the mechanism of toxicity for antibody-drug conjugates (ADCs) bearing non-cleavable and cleavable valine-citrulline (VC) linkers, which are known to have critically different efficacies toward different tumors^4, 12, 15, 16, 45^. We identified many known and novel genetic modifiers of ADC toxicity, and uncovered critical roles for a number of endolysosomal regulators in mediating the toxicity of ADCs. We identified C18orf8/RMC1 as a gene required for endosomal maturation and lysosomal delivery of ADC. Surprisingly, only a subset of the identified endolysosomal regulators are required for toxicity of ADCs with cleavable VC linkers. We showed that this is because lysosomal delivery is not required for payload cleavage from VC-linked ADCs, which contrasts with the prevailing wisdom in the field. Finally, our studies revealed a role for sialic acids in regulating ADC uptake, and we propose a novel combination therapy strategy that combines ADC with inhibition of sialic acid biosynthesis to increase ADC delivery. In proof-of-principle studies, this strategy was effective in increasing *in vitro* killing with both CD22- and Her2-targeting ADCs in multiple cancer cell lines, including lymphoma, breast, and gastric cancer.

Although ADCs with noncleavable linkers are more stable and thus elicit less off-target toxicity when used *in vivo*^2, 45^, they are not effective against all ADC targets, and some tumors show marked resistance to these agents^4, 16, 45^. Our work shows that noncleavable-linker ADCs show greater reliance on late endosomal trafficking regulators (RAB7, RMC1, WDR81, and WDR91) and lysosomal cathepsins (Cathepsins A, C, and S) for toxicity, whereas surprisingly ADCs with VC-cleavable linkers (designed to be cleaved by cathepsin B), do not. Indeed, payload release from VC-linked ADCs occurs within tens of minutes after ADC internalization, suggesting that resistance to noncleavable ADCs due to impaired lysosomal delivery may be circumvented by ADCs bearing cleavable VC linkers. These results may help explain why ADCs with cleavable linkers are efficacious in a broader range of target cancers^12, 16^ and remain toxic to cell lines resistant to noncleavable linker ADCs^46–49^. Importantly, these genes regulating late endosomal trafficking and processing of ADCs may serve as potential predictive biomarkers for resistance against noncleavable linker ADCs.

Our work also identified a number of new regulators of endolysosomal trafficking, as well as new roles for previously characterized genes. For example, we find that WDR81 and WDR91, genes previously shown to be involved in endosomal maturation and autophagy^29, 30, 50, 51^, are also selectively required for trafficking of ADCs to the lysosome. In addition, we demonstrate a new role for C18orf8/RMC1 in intracellular trafficking. This gene was recently identified to play a role in autophagy, and was found in a complex with CCZ1 and MON1^40^, which we also identify in our screen. Here, we show that RMC1 is required for proper endolysosomal trafficking and degradation of EGFR. Additionally, RMC1 mediates ADC delivery and toxicity in a RAB7A-dependent manner, which is consistent with the reported role of RMC1 in RAB7A activation during autophagic flux^40^.

Many of the hits identified in our study are required for ADC toxicity; therefore loss-of-function of these genes may be potential resistance mechanisms for ADCs used in the clinic. For example, decreased expression of SLC46A3, a lysosomal transporter we identified as critical for cytotoxicity of the anti-CD22 noncleavable ADC, has been shown to be a mechanism of innate and acquired resistance to other noncleavable ADCs targeting a wide range of antigenic targets in patient-derived xenografts and primary multiple myeloma bone marrow samples^22, 47, 52^. Moreover, a major mechanism of acquired resistance to ADCs shown in *in vitro* models is altered ADC trafficking and lysosomal function^47, 53, 54^, a signature that is also identified in our screens.

We found that inhibition of sialic acid biosynthesis renders cells more sensitive to ADCs due to an increased rate of ADC uptake in a wide range of cancer cell types. Previous studies have shown that altering cell surface sialic acid levels can influence receptor uptake through disrupting Siglec receptor clustering^55^, membrane fluidity^56^, or binding of cell surface glycans to the extracellular galectin lattice^57, 58^. Further research will be required to determine whether these mechanisms play a role in influencing the rate of ADC uptake.

Because hypersialylation in malignant cells has been shown to be a mechanism of immune evasion^59–61^, preclinical studies focusing on depleting tumor sialylation have developed several promising therapeutic strategies^62^. One such example is the fluorinated sialic acid analogue we used in our study, 3F-NeuAc. Our data show that combining 3F-NeuAc with either anti-CD22 ADCs or anti-Her2 ADC T-DM1 resulted in increased killing. Notably, 3F-NeuAc is effective in reducing global glycan sialylation in mice but has deleterious “on-target” effects such as impairing liver and kidney function ^63^. This systemic toxicity can be circumvented by intratumoral injections of 3F-NeuAc, resulting in decreased tumor sialylation levels^64^. Another possible means to implement this strategy would be to utilize targeted-glycan modification, in which sialidases conjugated to antibodies are recruited to target cells to remove cell surface sialic acids^65^. Our work suggests that combining ADCs with these and other existing methods to deplete sialylation in cancer cells may be a promising therapeutic strategy to increase ADC efficacy that warrants further exploration. Together, our studies reveal mechanisms underlying ADC trafficking and toxicity, provide insight for ADC design and a generalizable method to investigate the biology of diverse ADCs, and identify candidate combination strategies that may improve therapeutic efficacy.

## Online Methods

### Cell Culture

Ramos cells, SKBR3, ZR-75-1, and NCI-N87 cells were obtained from ATCC and grown in Roswell Park Memorial Institute (RPMI) 1640 Medium (11875093, Life Technologies) supplemented with 10% Fetal Bovine Serum (Fisher, Cat# SH30910), 2 mM L-glutamine (Fisher, Cat# SH3003401) and 1% penicillin-streptomycin (Fisher, Cat#SV30010). HeLa cells (ATCC) were grown in Dulbecco’s Modified Eagle’s Medium (Life Technologies, Cat# 11995073) supplemented with 10% FBS, 2 mM L-glutamine, and 1% penicillin-streptomycin. All cells were cultured at 37 °C with 5% CO^2^.

### Genome-wide CRISPR-Cas9 screens in Ramos cells

Our previously established genome-wide, 10sgRNA/gene CRISPR-deletion library^20^ was synthesized, cloned, and infected into Cas9 expressing Ramos cells as previously described^20^. Briefly, ∼300 million Ramos cells stably expressing SFFV-Cas9-BFP were infected with the CRISPR-knockout library at an MOI of 0.3-0.4. Cells expressing sgRNAs were selected for using puromycin (1 µg/mL) for 3 to 4 days such that >90% of cells were mCherry positive as measured by flow-cytometry. Selected cells are then allowed to recover and expand in puromycin-free media for up to 7 days. Deep sequencing was used to confirm sufficient sgRNA representation in the library.

For the screen, cells were split into two conditions, each in duplicate: an untreated control group and ADC-treated group (2 nM anti-CD22-Asp-PEG2-maytansine). Cells are incubated with 2 nM ADC for 48 hours for each round of treatment, after which the ADC was removed from media by pelleting cells and re-suspending in fresh ADC-free media. This treatment occurred four times over three weeks, with cells recovering to ∼90% viability after each round of selection. Over the course of the screen, cells were maintained at 500,000 cells/mL. At the end of the screen, 200 million cells were recovered from each condition and pelleted by centrifugation. Genomic DNA of each condition was extracted using Qiagen DNA Blood Maxi kit (Cat#51194). The sgRNA sequences were amplified and prepared for sequencing as previously described^21^. These libraries were then sequenced using an Illumina NextSeq with ∼40 million reads per condition; ∼200X coverage per library element). Analysis and comparison of guide composition of ADC treated versus untreated conditions were performed using casTLE as previously described^20, 21^. See Supplementary Table 1 for complete genome-wide screen result.

### Targeted screens in Ramos cells

The secondary screen library targets a total of 850 genes (10 sgRNA per gene): all genes that passed 10% FDR from the Ramos genome-wide screen (168 genes), genes previously implicated in endolysosomal trafficking, and genes found to localize to lysosomes in a proteomics study^34^, along with 1500 negative control sgRNAs (see Supplementary Table 5 for complete list of genes and sgRNA). The library oligos were synthesized by Agilent Technologies and cloned into pMCB320 using BstXI/BlpI overhangs after PCR amplification. The Cas9-expressing Ramos cells were lentivirally infected with the secondary screen library at an MOI of 0.3-0.4 as previously described. After puromycin selection (1µg/mL for 3 days) and expansion, cells were treated with either 2 nM anti-CD22-maytansine, 0.5 nM anti-CD22-VC-Maytansine, 0.2 nM free maytansine, or left untreated as control, each in duplicate at ∼3000X coverage per library element. Each treatment lasted 48 hours and was repeated four times over three weeks, with cells recovering to ∼90% viability after each round of selection. As with the genome-wide screen, cells were maintained at 500,000 cells/mL throughout the screen. At the end of the screen, 30 million cells were used for genomic extraction for each condition, sequenced (5-10 million reads per condition) and analyzed using casTLE. See Supplementary Table 2 for complete sublibrary screen results.

### Targeted screen for uncovering sensitizing hits

To uncover additional sensitizing hits, we used a library of ∼2000 genes targeting known drug targets, kinases and phosphatase in the human genome^20^. The library was lentivirally infected into Cas9-expressing Ramos cells at an MOI of 0.3-0.4 as previously described. After puromycin selection (1 µg/mL for 3 days) and expansion, cells were treated with either 0.5 nM anti-CD22 noncleavable linker ADC, 0.1 nM free maytansine, or left untreated, each in duplicate at ∼5000X coverage per library element. Each treatment lasted 24 hours, killing 20-30% of cells. Cells were allowed to recover to 90% survival after each treatment. The treatment was repeated three times over two weeks. Cells were maintained at 500,000 cells/mL throughout the screen. At the end of the screen, 50 million cells were used for genomic extraction for each condition and sequenced with 10 million reads per condition. Hits were identified using casTLE. See Supplementary Table 3 for complete screen result.

### Competitive growth assays in Ramos cells

To validate results of genome-wide and secondary screens, we infected Cas9-expressing Ramos cells with plasmids expressing both a single specific sgRNA and mCherry, as previously described^66^ (See Supplementary Information for sgRNA sequences). The infected cells were puromycin-selected for 3 days and allowed to recover for 2 days. Equal numbers of sgRNA/ mCherry-positive cells were cocultured with wildtype/ mCherry-negative cells, and were subsequently treated with different ADCs at the same concentration used in the screens. The percentage of mCherry-positive cells was quantified using a BD Accuri C6 flow cytometer after 72 hours. All results are normalized to ratio of control sgRNA/ mCherry-positive: wildtype/ mCherry-negative cells.

### Western Blotting

Cells were lysed for 30min at 4C using protein extraction buffer (300mM NaCl, 100mL Tris pH 8, 0.2mM EDTA, 0.1% Triton X-100, 10% glycerol) supplemented with 1X cOmplete protease inhibitor (Roche Cat#11697498001) and centrifuged to collect the supernatant lysate. Protein concentration was measured by DC Protein Assay Kit (BioRad Cat# 5000111EDU), separated on SDS-PAGE gels (Life Technologies Cat#NP0322BOX) and transferred to nitrocellulose membranes (BioRad Cat#1620146). The following primary antibodies were used: rabbit monoclonal anti-EGFR (1:2000, Abcam ab52894) and mouse monoclonal anti-GAPDH (1:10000, Fisher AM4300). Secondary antibodies used were: goat polyclonal anti-mouse IRDye 680RD (1:10000, Li-COR 925-68070) and goat-polyclonal anti-rabbit IRDye 800CW (1:10000, Li-COR 925-32211). Images were acquired using the Li-COR Odyssey CLx imaging system.

### Confocal microscopy

Cells were grown on glass coverslips or glass-well plates and were stained using standard immunocytochemistry techniques. Briefly, cells were fixed with 4% formaldehyde, permeabilized with 0.1% Triton X-100, blocked with 3% Bovine Serum Albumin, and stained with the following antibodies were used: mouse monoclonal anti-EEA1 (1:1000, BD 610457) and rabbit monoclonal anti-LAMP1 (1:1000, CST 9091S). Coverslips were mounted using VectaShield. Images were acquired using Nikon Eclipse microscope with a 100x oil immersion objective.

### Sample preparation and measurement of ADC catabolite by Liquid chromatography/ Mass spectrometry (LC/MS-MS)

For each condition, 10 million Ramos cell, or 5 million HeLa cells were plated in 10cm dishes and allowed to grow for 24 hours prior to addition of 10 nM anti-CD22-VC-maytansine or T-DM1. At appropriate time points, cells were collected (trypsinized in the case of HeLa cells) and pelleted. The cell pellets were washed 2x with PBS and extracted with 50% acetonitrile in water containing 10 nM MMAE as internal standard. The cell/acetonitrile mixture was vortexed for 30 seconds, and allowed to rest on ice for 1 minute, and this vortex / rest step was repeated two more times. The mixture was then left to shake at 4 °C for 15 minutes, incubated at −20 °C for 2 hours, and centrifuged at 10000g. The clarified supernatant was transferred to a new tube and dried under N^2^ gas flow. The dried extract was reconstituted in 100uL 50% acetonitrile in water prior to analysis by liquid chromatography-tandem mass spectroscopy (LC-MS/MS), which was performed using an Agilent 6470 triple quadrupole mass spectrometer with an Agilent Jet Stream (AJS) electrospray ionization source. Analytes were separated on a ZORBAX RRHD Extend C18 column (1.8 μm particle size x 2.1 mm diameter x 50 mm length) with by gradient elution at a flow of 0.6 mL/min. The mobile phases were A: 0.1% formic acid in water and B: 0.1% formic acid in acetonitrile. Gradient conditions were: isocratic at 95% A for 0.2 min, followed by a linear gradient from 5% to 95% B from 0.2 to 4 minutes, followed by an isocratic hold at 95% B for one minute (4.2-5.2min), followed by immediate return to 5% A (5.2-5.3 min). The total run time was 6 min. The ion source conditions were: gas temperature at 250°C, gas flow at 12l/min, nebulizer at 25psi, and sheath gas temperature and flow rate were at 300°C and 12l/min respectively.

Multiple-reaction monitoring (MRM) was used to monitor the analytes and internal standard. The mass transition monitored for quantification were m/z 749.4 → 547.2 for maytansinol payload released from VC linker, m/z 1103.6 → 547.2 for Lys-MCC-DM1, and m/z 718.5 → 686.5 for MMAE. For Lys-MCC-DM1 and MMAE, we confirmed that observed chromatographic peaks eluted at the same time as authentic standards. For the maytansinol payload released from the VC linker, no authentic standard was available. During methods development, we used additional MRMs m/z 749.4 → 485.2, 1103.6 → 485.2, and 718.5 → 152.1 to confirm the identity of the released maytansinol payload, Lys-MCC-DM1, and MMAE respectively. Peak detection and integration were performed with Agilent MassHunter software. For Fig. 3c, 6d, and Supplementary Fig 3d, 3e, and 6d, the obtained ion counts in biological samples were normalized to ion counts for the MMAE internal standard, and this ratio was normalized to cell number.

### ADC uptake assays in IncuCyte

For uptake assays with Ramos cells, cells were seeded at 10000 cells per well in poly-L-lysine (0.1% w/v) coated 96-well plate and allowed to adhere overnight. Anti-CD22 ADC were labeled with IncuCyte Human FabFluor-pH Red antibody labeling reagent (IncuCyte Cat#4722) by mixing ADC and FabFluor reagent at molar ratio of 1:3 in RPMI media and incubating for 15 min. The antibody-FabFluor mixture was added to cells to a final concentration of 4 µg/mL. The cell plate is then transferred into an IncuCyte s3 and images were obtained with a 20x objective every hour for 24 hours. Analysis were performed with IncuCyte s3 integrated software.

## Acknowledgements

We thank G. Hess, A. Li, P. Drake, M. Grey, A. Lu, and S. Pfeffer for assistance and discussions; A. Gupta for help with LC/MS-MS, M. Dubreuil and E. Jeng for comments on the manuscript. This work was funded by grants from NIH grants F31 GM126688-01A1 (C.K.T.), NIH 1DP2HD084069-01 (M.C.B.), and NIH R01 CA227942 (C.R.B.).

## Author Contributions

C.K.T. designed and performed experiments, analyzed data, and wrote the manuscript. R.M.B. generated ADC reagents. C.R.F. helped design and assist experiments performed with LC/MS-MS. D.W.M. assisted in screen data analysis. A.L. assisted in library cloning and screens. B.S. and C.R.B. helped design sialic acid inhibition experiments. M.C.B. and D.R. supervised the study.

## Competing interests

R.M.B. and D.R. are employees of Catalent Biologics. C.R.B. is a cofounder and Scientific Advisory Board member of Redwood Bioscience, which generated antibody-drug conjugates used in this work. Stanford University has filed a patent application based on the findings in this article.

---

**Supplementary Figure 1:**
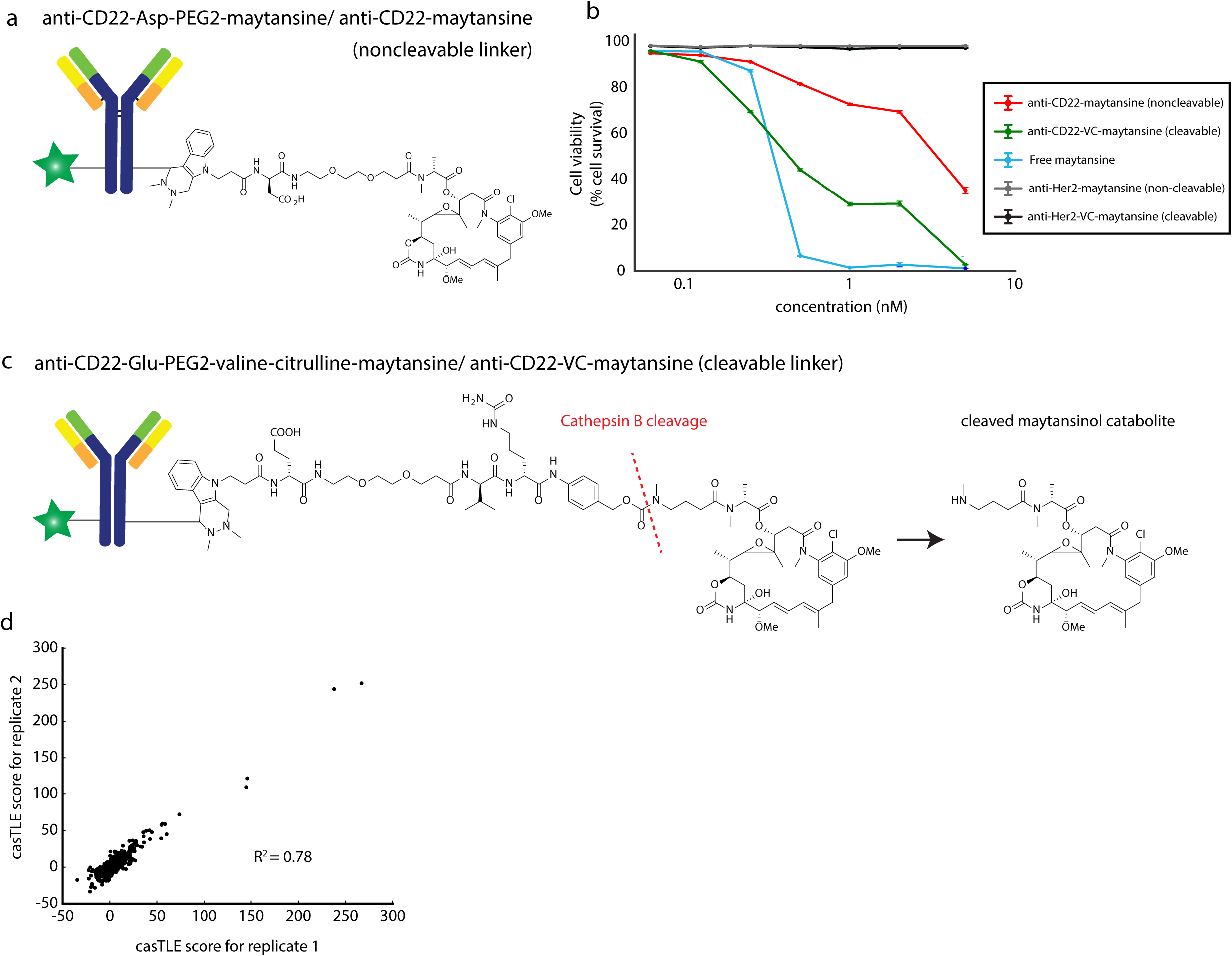
ADC structures and their in vitro effi cacy. a. Linker and payload structure of anti-CD22-maytansine (noncleavable linker) ADC. b. Cell viability of Ramos cells treated with indicated ADCs after 3 days. Viability was as determined by forward and side scatter live gating of Ramos cells (n=3; error bars, ± SEM) c. Linker and payload structure of anti-CD22-VC-maytansine (cleavable linker) ADC. Also depicted is the predicted cathepsin B cleavage site and the resulting catabolite. d. Replication plot of genome-wide screen in Fig. 1b.

**Supplementary Figure 2:**
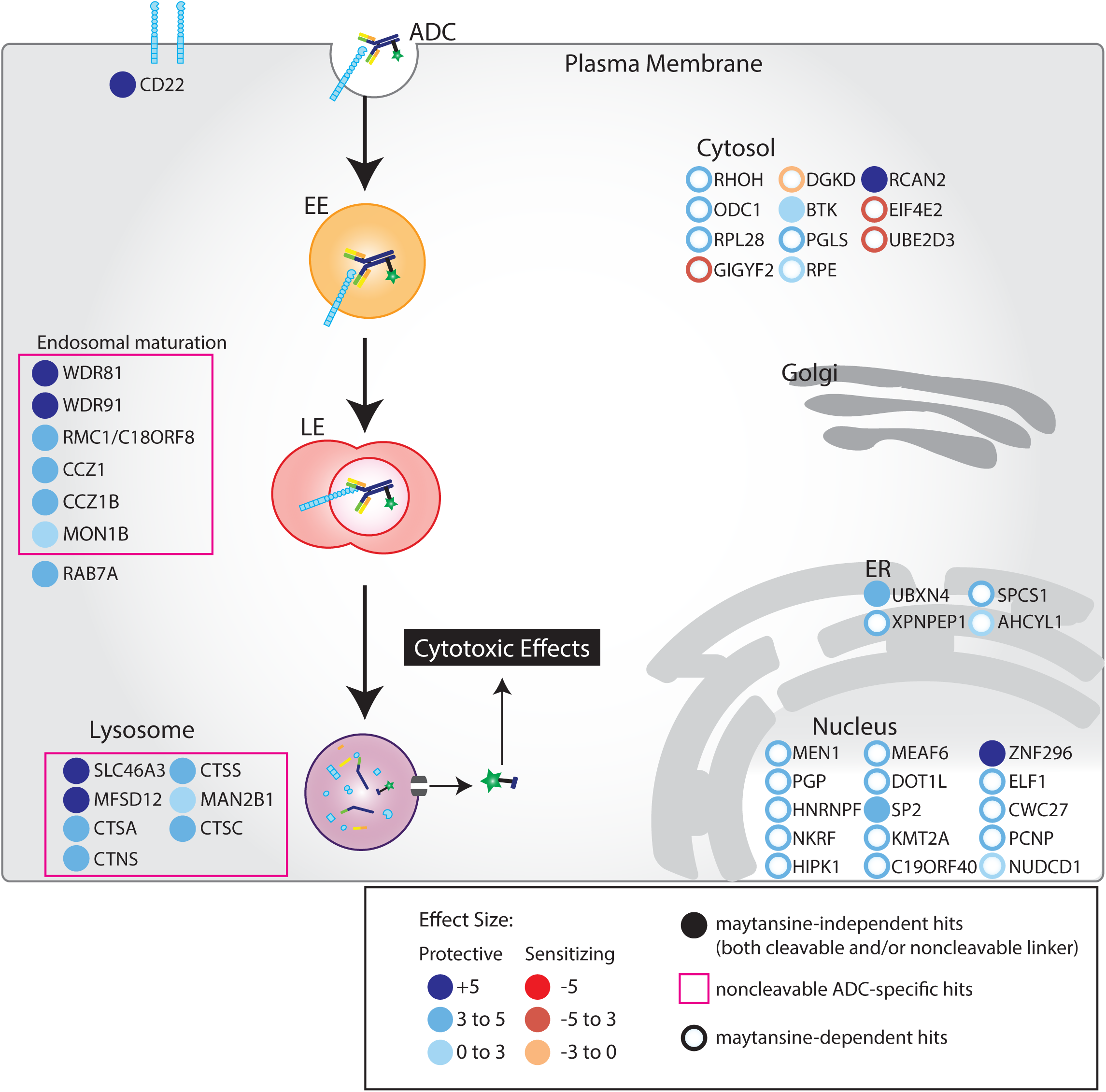
Summary of screen hits from anti-CD22-maytansine (noncleavable), anti-CD22-VC maytansine (cleavable), and free maytansine. Schematic for screen hits from anti-CD22-maytansine (noncleavable), anti-CD22-VC-maytansine (cleavable), and free maytansine. Depicted are the top 30 hits and selected endolysosomal genes, color coded by effect size (calculated by casTLE). Filled in circles represent maytansine-independent hits and empty circles represent maytansine-dependent hits. Hits unique to noncleavable linker ADC is denoted by pink box. Subcellular localization are indicated according to GO annotations.

**Supplementary Figure 3:**
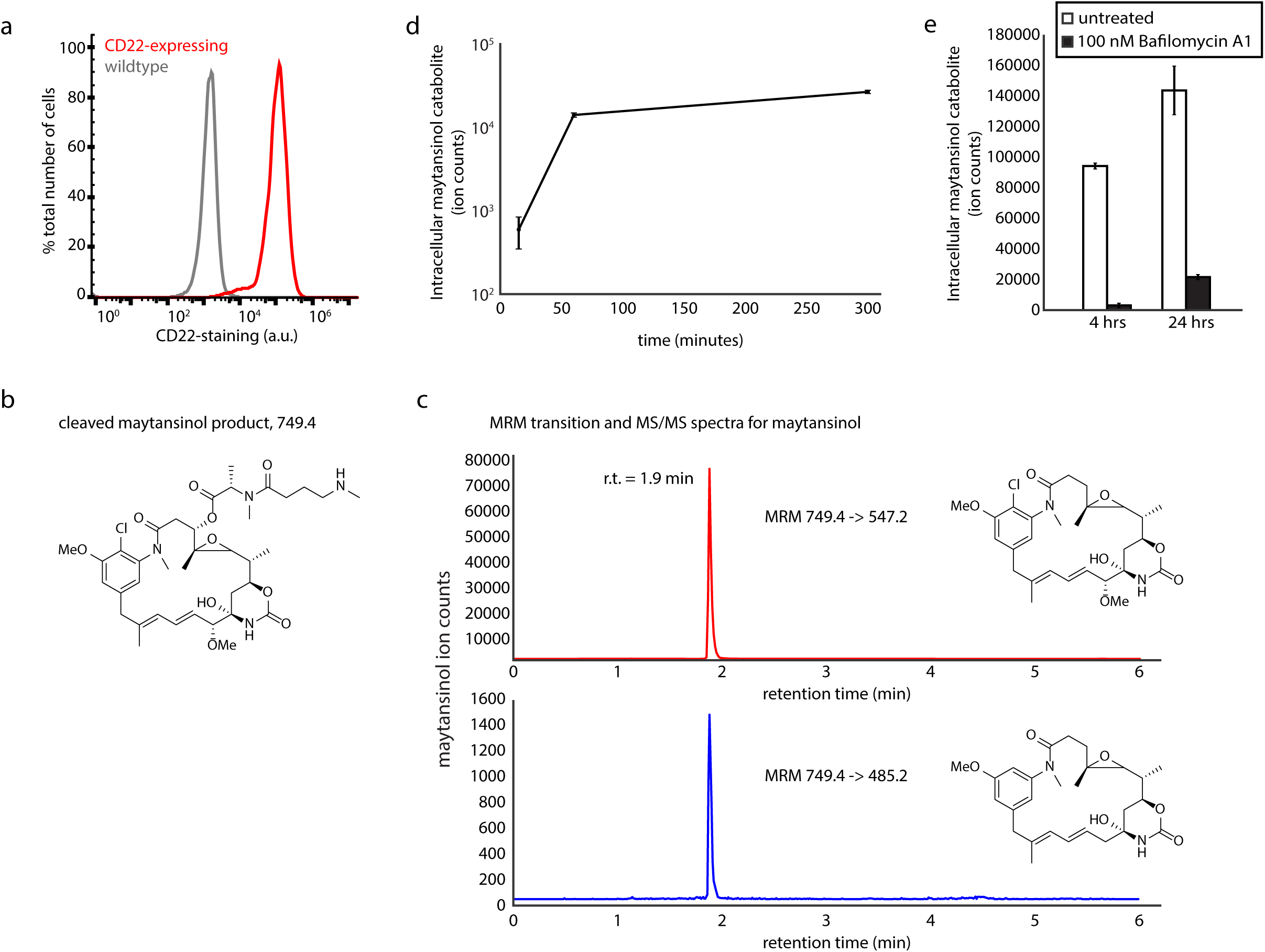
Detection of ADC catabolite by LC/MS-MS. a. Cell surface CD22-staining in HeLa cells stably expressing CD22, measured using anti-CD22-AF488 antibody followed by flow cytometry (n=3; error bars, ± SEM). b. Structure of cleaved maytansinol payload. c. MRM transition and MS/MS spectra for maytansinol payload. d. Maytansinol release from anti-CD22-VC-maytansine in CD22-expressing HeLa cells. Following anti-CD22-VC-maytansine addition, level of intracellular maytansinol catabolite was determined by LC/MS-MS at indicated time points; see methods for detailed extraction and detection protocols (n=3, normalized by cell number and internal standard MMAE; error bars, ± SEM). e. Maytansinol release from anti-CD22-VC-maytansine in CD22-expressing HeLa cells treated with Bafilomycin A1. Cells were treated with Bafilomycin A1 or left untreated for 20 hours. Cells were then incubated with anti-CD22-VC-maytansine for indicated times. Level of intracellular maytansinol catabolite was determined by LC/MS-MS; see methods for detailed extraction and detection protocol (n=3, normalized by cell number and internal standard MMAE; error bars, ± SEM).

**Supplementary Figure 4:**
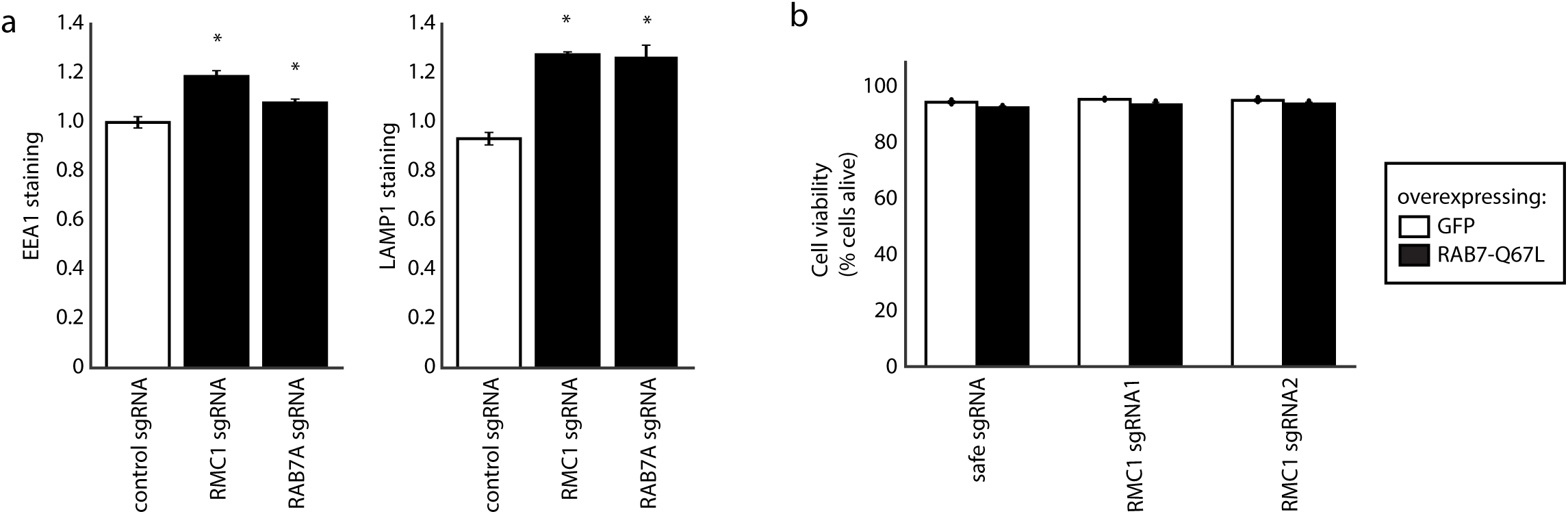
Depletion of RMC1 alters endosomal and lysosomal morphology. a. Total levels of EEA1 and LAMP1 in HeLa cells expressing the indicated CRISPRi sgRNA constructs. Cells were fixed, permeabilized, and stained with anti-EEA1 and anti-LAMP1 antibodies, followed by fluorescent secondary antibodies, and assayed by flow cytometry (n=3; Two-tailed t test, *P<0.05; error bars, ± SEM). b. Viability of Ramos cells overexpressing constitutively active RAB7A-Q67L. Cells were left untreated for three days and their viability, determined by forward and side scatter live gating, was measure by flow cytometry (n=3; error bars, ± SEM).

**Supplementary Figure 5:**
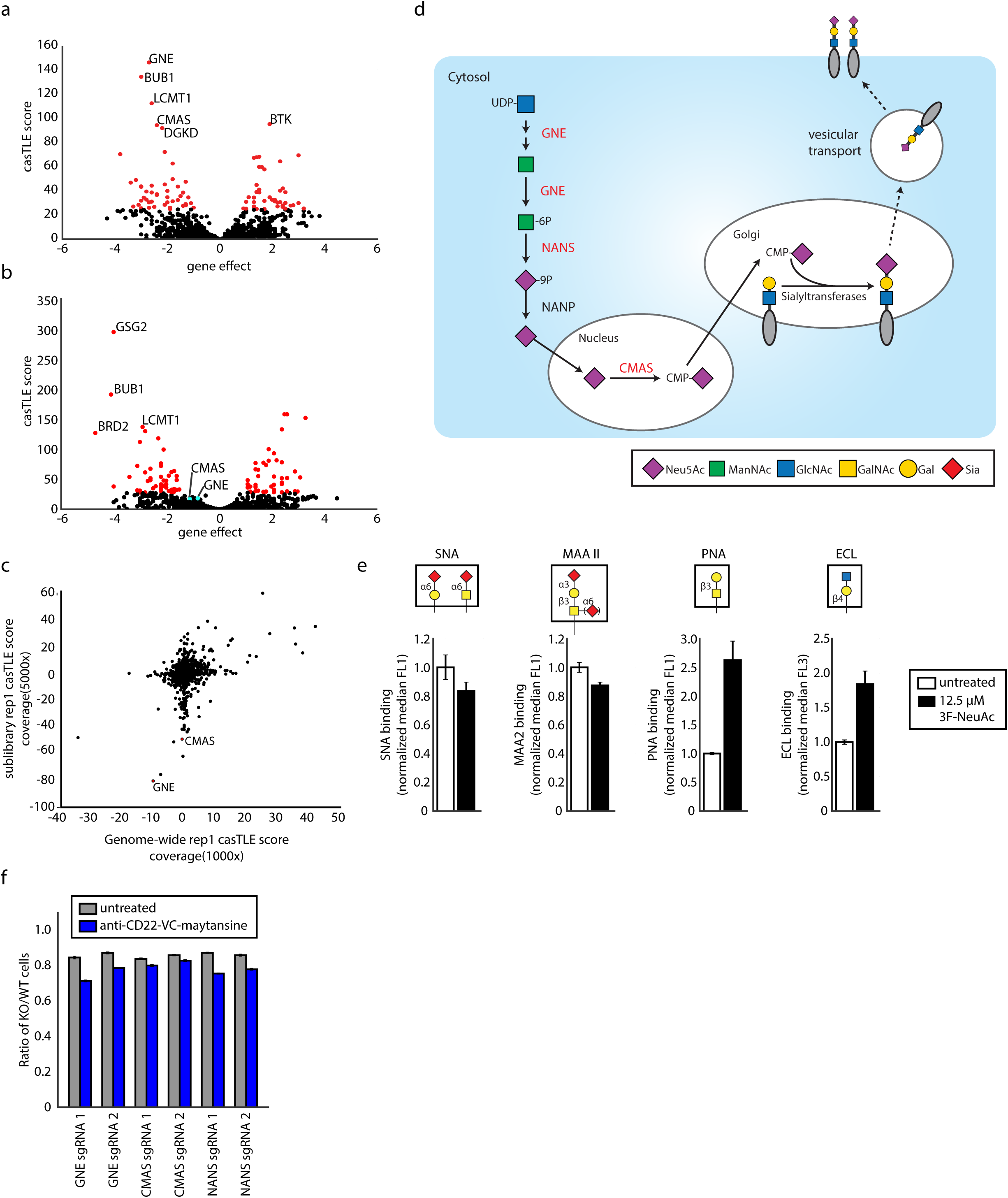
Targeted screen uncovers role of sialic acid biosynthesis in regulation of ADC toxicity. a. Volcano plot of all genes indicating effect and confidence scores for the drug target, kinases and phosphatases sublibrary anti-CD22-maytansine (noncleavable) screen. Effect and confidence scores determined by casTLE. b. Volcano plot of all genes indicating effect and confidence scores for the drug target, kinases and phosphatases sublibrary free maytansine screen. Effect and confidence scores determined by casTLE. c. Comparative analysis of results from genome-wide (1000x coverage, 2 nM anti-CD22-maytansine) and sublibrary (5000x coverage, 0.5 nM anti-CD22-maytansine) screens. Signed casTLE scores are reported. d. Schematic for sialic acid synthesis pathway. Genes in red are validated using competitive assay shown in Fig. 5c. e. Levels of cell surface sialic acid and“uncapped” glycans on Ramos cells treated with 12.5µM 3F-NeuAc as detected by the indicated lectin. Ramos cells were treated with 3F-NeuAc for 48 hours, washed and stained by indicated lectins. The binding preferences of each lectin are depicted (n=3; error bars, ± SEM). f. Competitive growth assays in Ramos cells using VC cleavable linker ADC. Cells expressing sgRNAs for KO of indicated genes (mCherry+) and control (mCherry-) were co-cultured in 1:1 ratio. Cells were either treated with anti-CD22-VC-maytansine or left untreated for three days. Resulting ratio of KO:control was determined using flow cytometry (n=3; error bars, ± SEM).

**Supplementary Figure 6:**
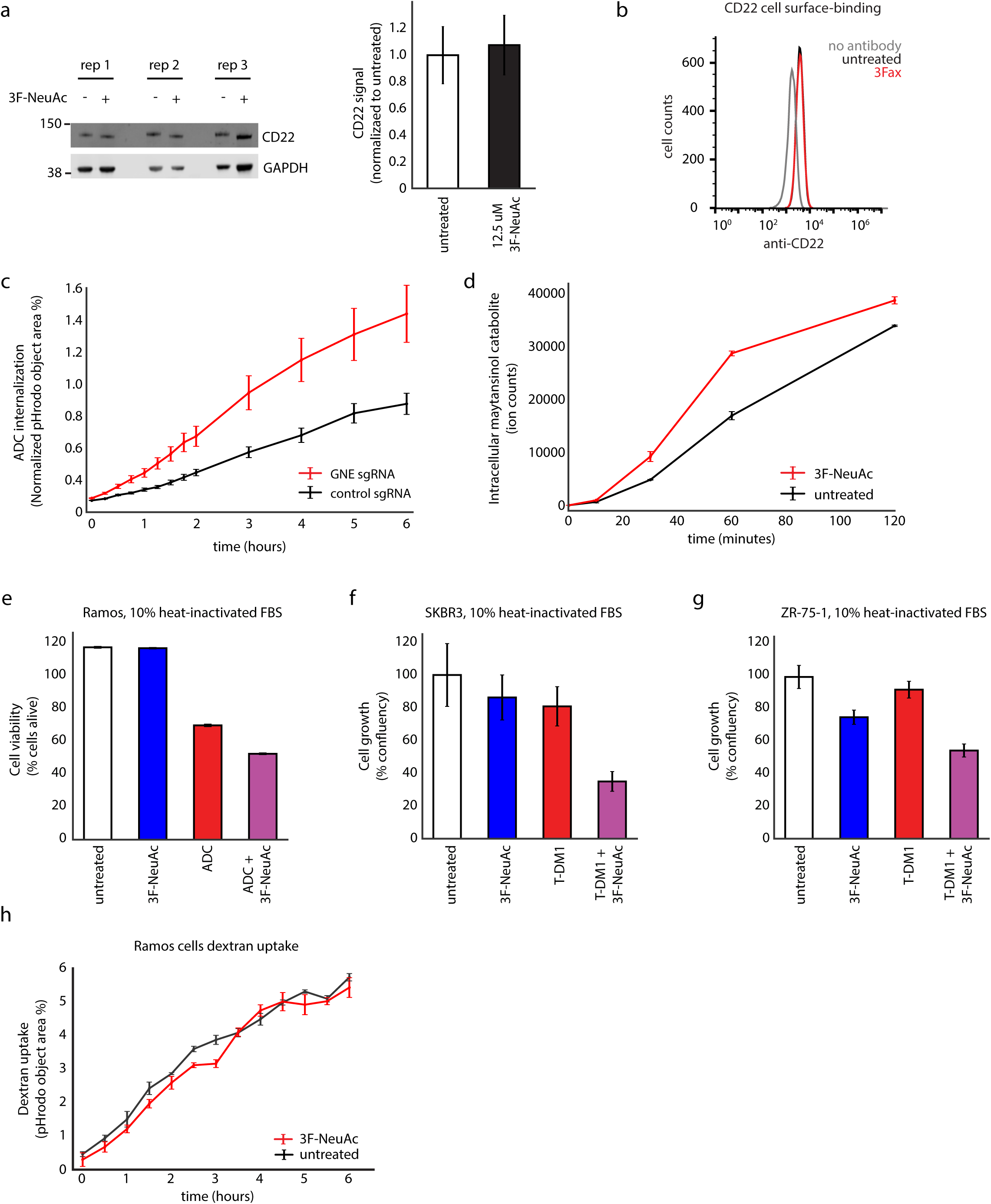
Increased sensitivity to ADC by inhibiting sialylation is not due to alteration of CD22 cell surface levels or enhanced complement-mediated cell killing. a. CD22 whole cell protein levels measured by Western blot in Ramos cells. Cells were treated with 12.5 µM 3F-NeuAc for 48 hours, washed, and lysed for protein extraction. Western blotting was used to detect levels of CD22 (n=3; error bars, ± SEM). b. CD22 cell surface staining measured by flow cytometry in Ramos cells. Cells were treated with 12.5 µM 3F-NeuAc for 48 hours, washed, stained with AF488-labeled anti-CD22 antibodies, and analyzed by flow cytometry (n=3; error bars, ± SEM). c. Internalization of pHrodo-labeled anti-CD22 ADC in Cas9-expressing Ramos cells with either GNE sgRNA (red) or control sgRNA (black). pHrodo signal was measured using IncuCyte s3 Live cells analysis system and normalized to cell area (n=3; error bars, ± SEM). d. Maytansinol payload release in 3F-NeuAc-treated Ramos cells. Cells were pre-treated with 3F-NeuAc 12.5 µM for 48 hours, followed by incubation with anti-CD22-VC-maytansine for indicated times. Level of intracellular maytansinol catabolite was determined by LC/MS-MS; see methods for detailed extraction and detection protocol (n=3, normalized by cell number and internal standard MMAE; error bars, ± SEM). e. Cell viability of Ramos cells treated with 3F-NeuAc, anti-CD22-maytansine, or combination of both in media supplemented with heat-inactivated fetal bovine serum, assayed by flow cytometry. Viability was as determined by forward and side scatter of live gating of Ramos cells (n=3; error bars, ± SEM). f-g. Cell growth (normalized confluency %) of SKBR3 (f) and ZR-75-1 (g) treated with 3F-NeuAc, T-DM1, or combination of both, in media supplemented with heat-inactivated fetal bovine serum. Cells were pre-treated with 100 µ M 3F-NeuAc for 48 hours, followed by incubation with 2 nM T-DM1. Confluency was determined by IncuCyte s3 Live cells analysis system and normalized to maximum confluency of the untreated condition at the end of the 5-day experiment (n=3; error bars, ± SEM). h. Internalization of pHrodo-labeled dextran in Ramos cells. pHrodo signal was measured using IncuCyte s3 Live cells analysis system and normalized to cell area (n=3; error bars, ± SEM) (n=3; error bars, ± SEM).

## Supplementary Table Legends

**Supplementary Table S1:** Genome-wide CRISPR screen results with anti-CD22-maytansine ADC in Ramos cells.

**Supplementary Table S2:** ADC/endolysosomal sublibrary screen results with anti-CD22-maytansine (noncleavable ADC), anti-CD22-VC-maytansine (cleavable ADC), and free maytansine in Ramos cells.

**Supplementary Table S3:** Drug target, kinases, and phosphatases sublibrary screen results with anti-CD22-maytansine (noncleavable ADC) and free maytansine in Ramos cells.

**Supplementary Table S4:** Count files for all screens in Ramos cells.

**Supplementary Table S5:** ADC/endolysosomal sublibrary sgRNA composition.

